# FireCloud, a scalable cloud-based platform for collaborative genome analysis: Strategies for reducing and controlling costs

**DOI:** 10.1101/209494

**Authors:** Chet Birger, Megan Hanna, Edward Salinas, Jason Neff, Gordon Saksena, Dimitri Livitz, Daniel Rosebrock, Chip Stewart, Ignaty Leshchiner, Alexander Baumann, Douglas Voet, Kristian Cibulskis, Eric Banks, Anthony Philippakis, Gad Getz

## Abstract

FireCloud, one of three NCI Cloud Pilots, is a collaborative genome analysis platform built on a cloud computing infrastructure. FireCloud aims to solve the many challenges presented by the increasingly large data sets and computing requirements employed in cancer research. However, cost uncertainty associated with cloud computing’s pay-as-you-go model is proving to be a barrier to adoption of cloud computing. In this paper we present guidelines for optimizing workflows to minimize cost and reduce latency. Our guidelines include: (i) dynamic disk sizing to efficiently utilize virtual disks; (ii) tuned provisioning of virtual machines (VMs) using a performance monitoring tool; (iii) taking advantage of steep price discounts of preemptible VMs; and (iv) utilizing the optimal parallelization of a task’s workload.

## Introduction

As sequencing costs plummet and the amount of genomic data generated soars [1], the extraction of biological insights from the data and the management of that data have emerged as significant challenges for researchers. The maturation and availability of large public data sets [2–5] from projects such as The Cancer Genome Atlas (TCGA) (<u>http://cancergenome.nih.gov/</u>), the International Cancer Genome Consortium (<u>http://www.icgc.org/</u>), and the Genotype-Tissue Expression (http://www.gtexportal.org/), stand to provide an invaluable opportunity to advance research, if the data can be made widely accessible and analyzable. At many institutions the challenges of reliably and cost-effectively running analyses over very large public and private datasets, and of storing these vast amounts of data, are pushing local compute and storage systems to their limits [1]. To overcome these big data challenges as a community, we must change the way research is conducted. We need to replace redundant and costly local infrastructure with shared resources [6, 7], provide infrastructure to reproducibly analyze and securely share data and results, make public data readily accessible to researchers, and transition to parallelizable and elastic compute and storage solutions. The National Cancer Institute’s (NCI) vision for the Genomic Data Commons (GDC) and Cloud Pilots[8] sets out to do that.

FireCloud (firecloud.org), one of the three NCI Cloud Pilots, addresses these challenges with the creation of a cancer genome analysis platform on a cloud computing environment that hosts all of TCGA data and is loaded with state-of-the-art tools and workflows. Using the elastic compute capacity of the Google Cloud Platform (GCP), FireCloud makes data readily accessible, facilitates collaboration and enables reproducible science by providing a robust, secure and scalable platform to the community at large. While FireCloud is a free service, compute and storage costs are charged to an attached Google account (for available credits, see Supplementary Text S4). FireCloud, like other emerging cloud-based solutions [9–11] provides analysts, tool developers, production managers and the broader biomedical research community with powerful tools and extensive resources to reliably compute across the increasingly large data sets.

FireCloud users manage and perform their work in workspaces. A FireCloud workspace is a computational sandbox that holds data, method configurations (instructions of how to run a workflow on the workspace’s data), and work history. Workspaces can be shared with other users to facilitate collaboration. Central to the workspace is the data model that consists of entities (e.g. samples, participants, pairs, and groups of them), the relationships among them, and attributes attached to these entities (attributes hold metadata that can be either explicit values or references to data files located on cloud storage). Workflows are executed on entities (or groups of entities) within a workspace, drawing inputs from attributes of the entities and writing outputs back as attributes, either as values or as references to files that are copied to the cloud storage (i.e., Google bucket) that is associated with the workspace. Method configurations contain the mappings of a workflow’s inputs and outputs to the specific attributes in the data model. Provenance is captured and managed for all analyses (executed workflows) run in a workspace.

FireCloud hosts predefined read-only workspaces that contain TCGA data; both Open (i.e. unrestricted data) and Controlled Access data (i.e. data that requires dbGaP authorization). FireCloud is a Trusted Partner for hosting TCGA data. FireCloud authenticates users through their Google (e.g., gmail) accounts. FireCloud manages authorization to TCGA controlled access data by requiring users to link their FireCloud account with their eRA Commons account through a second round of authentication with NIH iTrust. Users that have been granted access to TCGA controlled access data through dbGaP will then be able to enter FireCloud Workspaces with controlled-access data. We have recently expanded the available data sets to include image and proteomics data for TCGA samples and pediatric cancer data from the Therapeutically Applicable Research to Generate Effective Treatments (TARGET database; https://ocg.cancer.gov/programs/target). Users may clone these predefined workspaces to conduct their analyses on the data.

FireCloud is designed to support collaborative science with specific controls on who can access different resources. Creators of workspaces can grant access and rights to run jobs to other users. Similarly, the ability to access and run a workflow is controlled by its developer. The scalability and ability to securely share data, tools and results enables large-scale multi-institutional collaborations that were nearly impossible to perform otherwise. This collaborative environment will undoubtedly significantly accelerate cancer research, which is often delayed by cumbersome exchange of partially and non-homogeneously processed data.

The architecture of FireCloud is outlined in Figure 1 (a detailed description of the FireCloud system and its component services is available at our online documentation site, <u>www.firecloud.org)</u>. The FireCloud portal is designed for a large range of users, including researchers with backgrounds in biology, computational biology and bioinformatics, with a simple user interface that holds analysis workflows and tools. The site includes an online forum for the FireCloud user community.

**Figure. 1:**
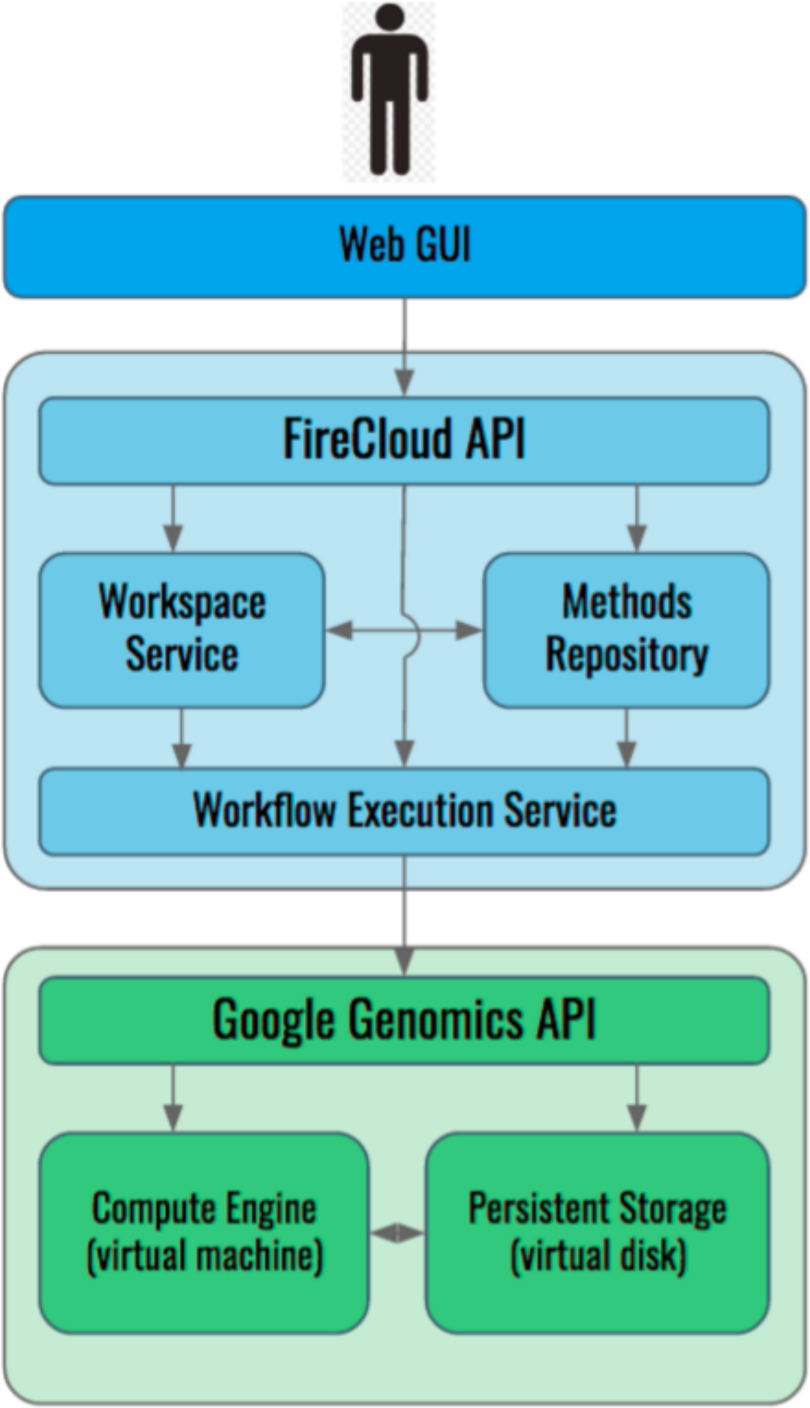
FireCloud schematic. FireCloud is a collaborative platform for genomic analysis that runs on the Google Cloud Platform. User interfaces are a Web GUI and a RESTful API for programmable access.

As the research community transitions from the traditional on-premises hardware solution (common in most research institutions) to cloud-based systems, numerous challenges have emerged. One of the most vexing, and perhaps a significant barrier to the research community’s adoption of cloud computing, is the move to the cloud’s pay-as-you-go pricing model. The cost of the traditional on-premises hardware model consists of a clear upfront cost for computer hardware (compute and storage) as well as an often ignored, continuous cost for technical maintenance and IT support. While controlling costs is straightforward and consistent with fixed budget grant funding, the compute power is limited by the available hardware and there is no way to handle surges in demand, which often occur in research projects. Hardware must be added regularly to keep up with the rapid escalation in data set sizes and related compute requirements. In contrast, cloud solutions do not require any capital outlay, and are more similar to a utility (pay-as-you-go) in that as much compute and storage as you need is generally available, and you are only charged for what you use, whether it is 10 minutes or 10 months. In our experience, capacity is not infinite, but the cloud can generally handle large surges in demand. While the scalable and elastic cloud environment addresses research’s spiky demand for compute power, and provides the ability to run large-scale analysis, there are still challenges such as the existence of caps and quotas on usage, difficulties in predicting and controlling costs, and the ease with which a user can incur cost overruns.

Moving to a cloud-computing environment brings with it a host of new issues and complexities for the users. For example, on most commercial clouds, upload of data to cloud storage is free, storage and compute are inexpensive, however, downloading data is relatively more expensive. This changes the way users need to think about and manage their data. Data that is readily retrieved from the source (for example, the GDC) can be deleted after the analysis is complete, leaving only the smaller (less expensive) analysis results files that can be downloaded, if needed. In addition to adjustments to data management, users will need to become familiar with the new cost model and the types of charges that can be accrued in running analysis on the cloud: hourly rates for the different VM types and associated virtual disk, costs for cloud storage options (ranging from $0.007 to $0.26 per GB/month), and data egress costs which vary by zone and volume. It is also important to understand where cost savings are available. For example, Google offers volume discounts for high-volume users, and also makes Preemptible VMs readily available at a steep discount (~80% discount). All major cloud providers offer similar pricing and discounts (AWS offers Spot Market at similar saving as Google Preemptibles).

Given that FireCloud enables the running of large-scale analysis on hundreds, or even hundreds of thousands of entities with the launch of a single command, it is possible to accumulate significant charges very quickly. For example, running the mutation calling workflow used in our testing for this paper (5 tasks, plus one 25-way scattered task; an overall of 30 tasks) across our test set of 100 TCGA Breast Cancer patients (100 patient-pairs of tumor and normal WES = 200 BAMs) requires storage for 4Tb (200 x ~18Gb = 3.6Tb) and requires 3,000 VMs (100 patient-pairs x 30 tasks = 3000) and 3,000 attached virtual disks. Time to process the 100 patient-pairs (wall clock time) is the time of the longest running patient tumor/normal pair (between 1 and 4 hours), whereas total (non-concurrent) time is closer to 1,000 hours. Currently there are only 2 caps on costs, a 7-day runtime limit on VMs, and a 24-hour limit on preemptible (see below) VMs, leaving much room for cost overruns. Although running on the cloud can decrease overall costs for the typical user, the difficulties of estimating and managing costs may deter researchers from migrating to the cloud. From the overall NCI perspective, storing the data in a small number of accessible cloud environments, rather than the current practice of storing many copies on different local systems, will provide substantial cost savings. Therefore, detailed understanding, control and management of costs in cloud-environments are key for enabling greater adoption of this new model.

To address these challenges, we have undertaken an evaluation and optimization effort with the goal of exploring different cost-performance tradeoffs and developing guidelines to assist FireCloud researchers, analysts and workflow developers in optimizing their workflows for cost as well as latency.

## Methods

For the purposes of this paper, we chose to analyze and optimize the performance of a version of the somatic mutation calling workflow available in FireCloud (called MutationCalling_MuTect). The test data is comprised of TCGA Breast Cancer (BRCA) data, both whole-exome sequencing (WES) and whole-genome sequencing (WGS) data (i.e. BAM files [https://github.com/samtools/hts-specs]). This workflow runs on a patient-matched tumor/normal pair, or a set of pairs, and consists of 6 separate tasks: (i)Prepare, (ii) MuTectSNV, (iii) MuTectFC, (iv) GatherAndOncotate, (v) VariantEffectPrediction and (vi) Report (see Supplementary Text S1 for more details). MuTectSNV (running the MuTect [12] somatic mutation caller), the most resource-expensive of these, is configured to be scattered 25 ways (i.e., the genomic coordinate space is split into 25 equally-sized intervals which are analyzed in parallel), and the GatherAndOncotate task merges the results.

Currently Google Cloud Platform does not report costs at the job level. However, we needed job-level cost reporting to optimize workflow and task costs and therefore found it necessary to develop our own cost estimation tools. For this analysis, we estimated costs by running a collection of tools (Supplementary Text S2) designed to gather job information (including job identifier, job runtime, virtual machine type, virtual disk size and type) and estimate runtime costs based on Google Cloud Platform pricing information.

In order to analyze the performance and cost of a workflow in FireCloud, one needs to understand how workflows are defined and executed. FireCloud workflows are comprised of one or more executable tasks stitched together into computational pipelines. Workflows are described in the Workflow Description Language (WDL) (https://software.broadinstitute.org/wdl) which is a domain-specific, human-readable language for describing workflows and their component tasks. Task definitions include specifications of their inputs, outputs and runtime attributes (e.g., Docker image, CPU, disk and memory requirements). Workflow definitions include specifications of workflow-level inputs and outputs, mappings of those inputs and outputs to task inputs and outputs, and inter-task “wiring” (declaring how the outputs of upstream tasks feed into the inputs of downstream tasks). Each executable task is run within a Docker container (<u>https://www.docker.com/)</u> hosted by a virtual machine running on the Google Cloud. The Docker container packages all of the code and system environment required to run a task as a single executable unit, enabling portability of task applications onto virtual machines running anywhere (either on the cloud or on a developer’s laptop). Docker images are snapshots of Docker containers, and are typically stored in Docker image repositories such as Docker Hub (https://hub.docker.com).

FireCloud uses GCP to run its workflows (Supplementary Figure S1). When a workflow is launched, FireCloud submits each of the workflow’s tasks to the Google Genomics API Service (v1alpha2 REST API, <u>https://cloud.google.com/genomics/reference/rest/</u>) for execution when the task’s inputs are available (inputs may be drawn from both the workflow’s inputs and upstream tasks’ outputs). The cost of a workflow is the total cost of all jobs (executed instances of a task) run in a workflow. The cost of a job is the sum of its charges for the virtual machine it runs on and the virtual disks attached to that machine. All VM instances are charged for a minimum of 10 minutes, and after 10 minutes, they are charged in 1-minute increments, rounded up to the nearest minute. Disk charges are prorated based on a granularity of seconds, with rates per Gigabytes of provisioned storage per month.

## Results

As somatic mutation calling is an essential part in the analysis of cancer samples and is typically run across a large number of tumor/normal pairs (in particular in large-scale sequencing projects), cost savings is our primary consideration in optimizing the MutationCalling_MuTect workflow. In some applications, for example in clinical settings, minimizing latency is of higher priority. Following are the steps we have taken to achieve cost savings with some considerations for latency reduction.

### Use dynamic disk-sizing to provision persistent storage (virtual disk)

In testing our mutation calling workflows on large cohorts, we found a low frequency of failures caused by insufficient disk space. Input files (e.g., BAM files) residing on cloud storage need to be copied onto a VM’s attached virtual disk (i.e., localized, which does not incur a charge) in order for a job to process the files. We had been statically provisioning the size of the attached virtual disk, and upon examination, saw that the failures were from the few outsized BAMs in the cohort. While our first impulse was to increase the statically specified, one-size-fits-all disk size, we realized that the incorporation of dynamic disk-sizing into our Prepare task, in addition to increasing fault tolerance, had cost saving implications. The workflow’s Prepare task now receives as input the sizes of all workflow input files and calculates the virtual disk size requirements for each of the down-stream tasks, thus ensuring that there are no oversized (and thus wasted) disks or undersized disks leading to failures.

### Tune provisioning of VMs with aid of a performance-monitoring tool

We struggled with higher-than-expected costs as we ran our mutation calling workflow on increasingly larger data sets, and we recognized the value of measuring the resource utilizations of each task in the workflow, especially MuTect, the most time consuming and thus costly task in our workflow. Accordingly, we incorporated dstat, (http://dag.wiee.rs/home-made/dstat/) a tool for runtime monitoring of utilization of CPU, memory and disk read and write, into each of the workflow’s tasks. Dstat-graph (http://lamada.eu/dstat-graph/) a companion tool, can present the results of dstat’s output logs in useful graphs that are easily interpretable.

Our examination of the dstat output (Figure 2) of the MuTect jobs showed that the CPU utilization, except for a few brief spikes, was just under 50%. We determined that FireCloud was requesting as its default VM type (n1-standard-2) Google Compute Engine’s standard machine type with 2 virtual CPUs and 7.5 GB of memory, but MuTect is a single-threaded application that can only utilize a single CPU per application instance. By adding runtime attributes that directed FireCloud to request a standard machine type with 1 virtual CPU and 3.75 GB of memory, excess CPU capacity was eliminated and the compute costs associated with the MuTect task were halved with no impact on task latency. Prior to this change VM costs for running MuTect on a single WES pair averaged $0.86; after moving to the single-CPU machine type, MuTect costs averaged $0.43 per WES pair (Table 1).

**Figure. 2:**
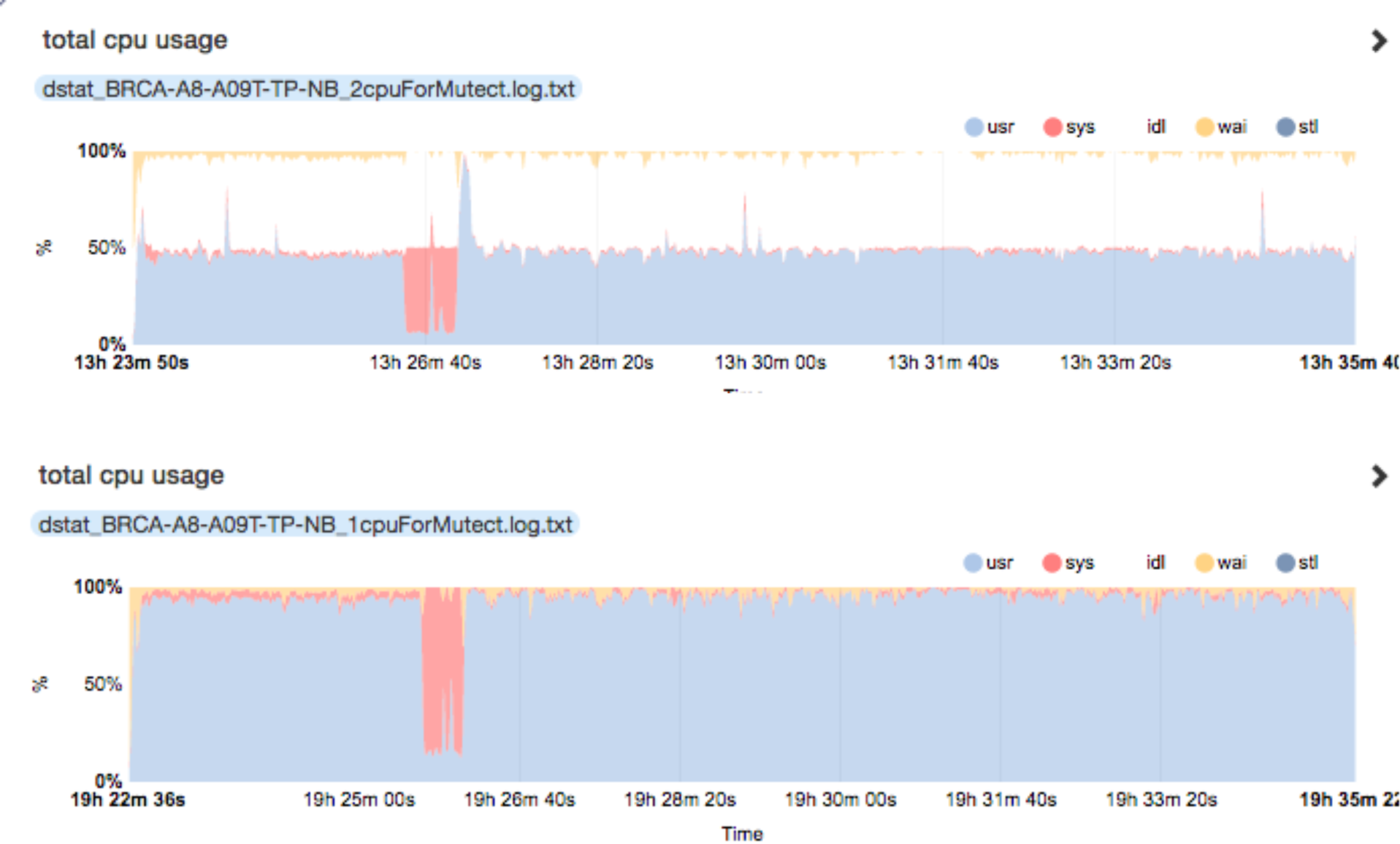
CPU utilization plot. Plot generated by applying dstat-graph (<u>http://lamada.eu/dstat-graph/</u>) on logs from the dstat resource-monitoring tool. MuTect task utilized only 50% of its VM’s total CPU capacity (top). FireCloud, by default, was requesting the standard virtual machine with 2 virtual CPUs and 7.5 GB of memory. MuTect is a single-threaded application that uses only a single CPU per instance. Requesting a virtual machine with 1 virtual CPU and 3.75 GB of memory, excess CPU capacity was eliminated (bottom) and the compute costs associated with the MuTect task were halved with no impact on task latency.

**Table 1.** Costs for running WES and WGS on Preemptible Machines. The mutation calling workflow was run on both WES and WGS data sets. For each data set, average cost per tumor/normal pair is reported for running MuTect on full price VMs and preemptible VMs. Costs are broken down into charges for the VM (CPU) and the virtual disk attached to the VM. Using preemptible VMs provided significant savings for both WES and WGS samples, but the savings were more pronounced for the WES cases, which had shorter processing times, and thus less chance of preemption.

| VM type | BAMs | Average Costs per pair |  |  | Average Latency per pair |  | Summary |  |
| --- | --- | --- | --- | --- | --- | --- | --- | --- |
| MuTect run over WES and WGS | Average size | CPU | Disk | Total VM | Total hours | Wallclock hours | Preemption rate | Savings |
| *WES Full price VM (95 pairs)* | *18Gb* | \$0.43 | \$0.04 | \$0.47 | 8.56 | 0.39 | | |
| *WES Preemptible VM (95 pairs)* | | \$0.09 | \$0.04 | \$0.13 | 8.64 | 0.38 | **0.04** | **73%** |
| *WGS Full price VM (9 pairs)* | *124Gb* | \$5.55 | \$3.53 | \$9.09 | 111.09 | 5.18 | | |
| *WGS Preemptible VM (9 pairs)* | | \$1.24 | \$3.83 | \$5.02 | 120.44 | 5.07 | **0.37** | **48%** |

### Use of preemptible virtual machines

FireCloud users can take advantage of the Google Cloud’s Preemptible VMs and potentially realize substantial reduction in workflow compute costs. Preemptible VM instances are excess compute capacity that Google makes available at a steep discount (up to 80% less expensive than normal instances). However, Google may terminate (preempt) these instances if it needs those resources for other uses and will always terminate preemptible instances after they run for 24 hours. Users are charged for preemptible VMs, regardless of whether or not the VMs are preempted. (Users are not charged if preemption occurs within the first 10 minutes of a job.)

One of the benefits of cloud computing is the seemingly infinite, ‘elastic’ compute; however, in running large-scale analysis, it is not uncommon to encounter congestion on the cloud platform. Furthermore, cloud-based platforms are not designed to provide 100% reliability. A goal of analysis platforms like FireCloud, therefore, are to create fault-tolerant computational services for users on top of cloud-based infrastructures.

This relieves analysts from building error recovery logic into their workflows and operations, and provides resilience in the presence of transient errors.

FireCloud’s workflow execution service (called Cromwell) is designed to be fault-tolerant to preemptions. It employs a simple retry policy when jobs running on preemptible VMs fail due to preemption: a task runtime parameter (defined in the WDL) indicates the maximum number of attempts Cromwell can make to run a task on a preemptible VM before running it on a full price non-preemptible machine. (If this parameter is set to 0, its default value, Cromwell makes no attempts to run the task on a preemptible VM and immediately runs it on a normal VM instance.)

While the cost savings achieved by running jobs on preemptible VMs can be considerable (Table 1), those savings are subject to the varying workload on the GCP. The probability that the Cloud Platform will terminate a preemptible VM instance, while generally low, will vary, and can spike during times of congestion within the Cloud.

We developed a simple stochastic model (Supplementary Text S3) for the compute costs (and latency) of tasks configured to run on preemptible VMs. Employing the model, we derived a formula for the expected cost ratio: the ratio of the expected cost of running a task on preemptible VMs and the cost of running the same task on a non-preemptible VM. We conducted thousands of runs of tasks with known execution times and calibrated our model with data collected from those runs. Having observed preemption rates within 10-minute intervals no greater than 5%, we set our simple model’s constant-valued hazard function λ(t) = λ to be ~0.3, where the unit of time is hours. With this calibrated hazard function value, our model led to the following guidelines for use of preemptibles:

1. <u>For jobs with (un-preempted) runtimes up to 8 hours in length, we recommend employing preemptible VMs.</u>
2. <u>For jobs with runtimes less than 4 hours, use preemptible VMs with N=5, where</u> N is the runtime parameter specifying the maximum number of attempts Cromwell makes before running the job on a normal VM
3. <u>For jobs with runtimes between 4 and 8 hours, use preemptible VMs with N = 2.</u>

Note that these guidelines are based on the preemption rates we observed on the Google Cloud Platform when our experiments were conducted. Those rates are subject to change.

### Leveraging multiprocessing to reduce task latencies in FireCloud workflows

Workflow latency (e.g., “wall clock runtime” includes both processing time and the time it takes to copy files to the virtual disk) can be reduced by parallelizing the processing activities required by the workflow. While inter-task dependencies constrain the level of parallelism amongst a workflow’s tasks, workflow developers can distribute an individual task’s workload across multiple instances of that task to reduce workflow latency. For many tasks, input data can be divided into smaller slices that can be processed in parallel. After all slices have been processed, the outputs can be assembled into a single output.

Currently there are two general approaches for leveraging multiprocessing to reduce task latencies in FireCloud workflows. The first is to employ WDL’s scatter clause to scatter an individual task’s workload across multiple instances of that task, each running on its own VM instance (each with an attached virtual disk) on the cloud; we will refer to this as an N-way scatter. The second is to run multiple instances of the same task on a single, large multi-core VM (with a single attached virtual disk) capable of running multiple processes in parallel; we will refer to this as M-way multiprocessing. Both approaches yield similar savings in overall task latency. In both cases, the latency is that of the longest running process. The question becomes which of the two is the most cost effective.

There are multiple factors impacting the relative costs of the two approaches: (i) the relative pricing of the small VM used in N-way scattering and the large VM used in M-way multiprocessing; (ii) the relative utilizations of the available CPU capacity for each of the two approaches; and (iii) the costs of persistent storage (i.e., attached disk). The majority of our somatic mutation calling workflow’s workload, and thus cost, is attributable to the running of the MuTect task. We compared the costs of scattering MuTect across 25 single-CPU virtual machines (N-way scatter, N=25) vs. the cost of running 32 instances of MuTect on a single 32-CPU virtual machine (M-way multiprocessing, M=32). We assessed the impact each of the above factors had on the relative costs of the two approaches. We ran the mutation calling workflow on both WES and WGS data sets.

#### Machine Pricing

Because Google’s per-CPU pricing ($0.04 per CPU core hour) is the same for all standard machine types, as long as standard machine types are used in both parallelization approaches, VM pricing differences alone cannot introduce a cost differential (32-CPU machine is 32 times more costly than a 1-CPU machine, but it can also process 32 times the workload). However, if the adoption of M-way multiprocessing requires the use of a “high-memory” machine type, the price differential (the per CPU pricing for a high-memory machine is $0.063 per CPU core hour) will make M-way multiprocessing more expensive than N-way scatter.

#### Utilizations of the available CPU capacity

Idle VM CPU capacity on a running VM incurs wasted charges, increasing the cost of running a workflow on the cloud. We encountered this when employing M-way multiprocessing for the MuTect task in our somatic mutation calling workflow. The reads were divided across 32 equal-size genomic intervals, and each interval had a single MuTect process dedicated to it. Because the task loads are not equal despite the evenly sized genomic intervals, the processes ended at different times, with the dstat logs and graphs revealing a long “tail” where all 32 of the VM’s CPUs were not fully utilized. This inefficiency is avoided with N-way scatter because VMs are terminated as soon their respective processes are completed. This was borne out in cost comparisons of N-way scatter and M-way multiprocessing (Figure 3). One way to remedy this issue and still employ M-way multiprocessing on the large 32-CPU VM is to divide a BAM’s reads across a larger number of genomic intervals. A single dedicated MuTect process will still process each interval, but as each process completes, a new MuTect process is launched to process the next genomic interval. The VM’s 32 CPUs will be kept busy as long as there are additional intervals to be processed. This will reduce the length of the tail, but not fully eliminate it (supplementary Figure S2).

**Figure. 3:**
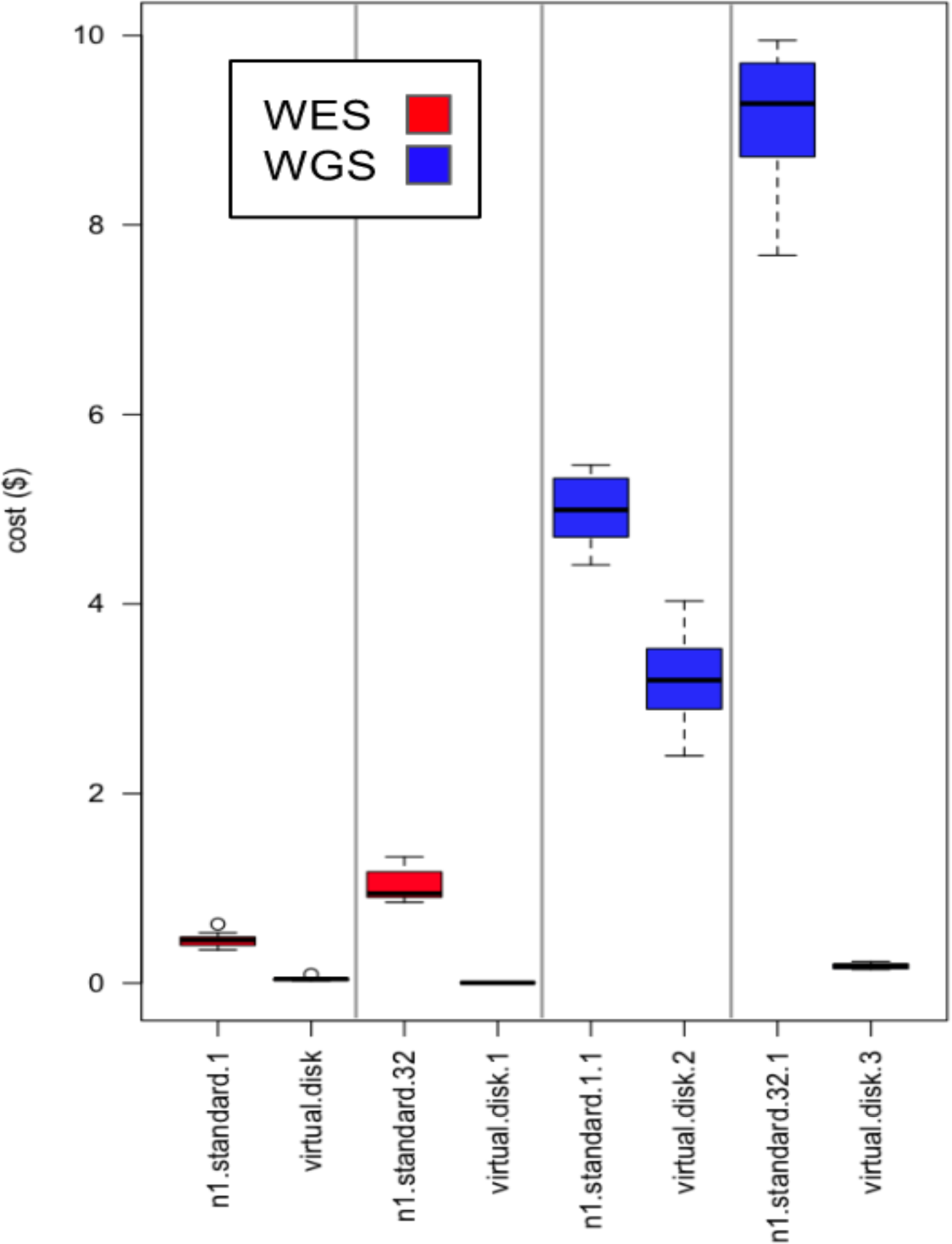
Comparing costs of running N-way scatter and M-way multiprocessing. Cost for the VMs and virtual disks are plotted for N-way scatter and M-way multiprocessing of the Mutect task. The N-way scatter runs 25 instances of the Mutect task, each on its own dedicated single-cpu n1.standard.1 VM. The M-way multiprocessing runs 32 instances of the Mutect application on a single large 32-cpu n1.standard.32 VM. We display box plots for both whole exome (red) and whole genome analyses (blue). For whole exome analyses, the N-way scatter is less costly due to the immediate termination of VMs once their processing of the task load is complete. As file sizes, and thus required disk sizes, grow, N-way scatter’s cost advantages shrink. For high-coverage whole genome files, M-way multiprocessing is more cost effective.

#### Costs of persistent storage (i.e., attached virtual disk)

If the persistent storage requirements for a single VM in the N-way scatter is 1/N that of the single VM in M-way multiprocessing, then the two approaches will have similar persistent storage costs. In our current system and design of the workflow, however, the persistent storage requirement of a single small VM is identical to that of the large VM because each VM instance requires a complete copy of both tumor and normal BAMs, despite the fact it is only operating on a slice of those BAMs. Thus, the aggregate cost of persistent storage for the N-way scatter is N times that of M-way multiprocessing, making the persistent storage costs of the N-way scatter significantly greater (by a factor of N) than the storage costs for M-way multiprocessing. For WES analysis (with typical BAM file sizes in the range of 15-25 GB) VM costs outweigh the costs of persistent storage attached to VMs (9% of the total costs are attributable to storage); even with the factor of N difference in storage costs between the two parallelization approaches, we found that N-way scatter’s larger persistent storage costs did not tip the cost-efficiency balance towards M-way multiprocessing. For WGS analysis (with typical BAM file sizes often exceeding 150 GB) persistent storage costs in an N-way scatter accounted for a much larger percentage of overall costs (at least 40%), making M-way multiprocessing at least as cost efficient at N-way scattering, and definitely less costly when working with high-coverage WGS BAMS (300 GB or larger).

Regardless of whether one chooses N-way scatter or M-way multiprocessing, we recommend employing preemptible VMs, subject to the preemptible guidelines we presented earlier. Note, however, that the preemption of the large VM used in M-way multiprocessing leads to a greater amount of work being lost and can lead to costs greater than those of running on non-preemptible VMs. The risks of this can be limited by keeping the preemptible parameter low (e.g., N ≤ 2).

## Discussion

For this paper, we set out to optimize a somatic mutation-calling workflow primarily for cost, but also for latency. From this effort, we gained insights into the sources that contribute to the cost and were then able to develop generalizable approaches that are applicable to other workflows. By increasing the visibility of cost contributors and providing approaches for managing them, we hope to help lower the barriers to the adoption of FireCloud and cloud computing in general.

General guidelines for optimizing workflows follow: i) use dynamic disk sizing to accurately size virtual disk; ii) tune provisioning of task VMs with aid of a performance monitoring tool like dstat to eliminate wasted CPU and memory resources; iii) take advantage of steep price discounts of preemtible VMs; iv) utilize the optimal parallelization of a task’s workload, either across multiple VMs or multiple processes within a single large VM. With the exception of the recommendations for use of Google’s Preemtible machines, our guidelines are generalizable to other commercial clouds.

We have optimized our publicly available best practice workflows in FireCloud (see Supplementary Text S5) in accordance with the guidelines above for the use of analysts and researchers. These workflows are publicly available for the FireCloud user community in the Method Repository. For pipeline developers, we have created a workspace (see Supplementary Text S5) that contains the methods that were used for the work on this paper. We recommend using these workspace methods as models for implementing the guidelines in this paper. To estimate workflow costs, we recommend running the optimized workflow on a small subset of data to capture costs first (charges can be viewed at *console.developers.google.com*). Users can then infer the costs of running the workflow on the larger dataset.

We believe that a cloud provider agnostic system is in the best interests of the community, however, for expediency we built FireCloud leveraging as many Google services as needed. We expect that future versions of FireCloud will be enabled to utilize other commercial clouds.

We anticipate that future technical innovations will further reduce the cost of running computational workflows on the cloud. For example, data streaming from cloud storage will avoid data localization and the resulting costs for virtual disk storage. In addition, we expect the development of useful tools to manage costs; for example, GCP or third party tools for more granular cost reporting, or real-time reporting of charges (metering). As new technologies for distributed computing are emerging, Spark (http://spark.apache.org/) for example, they will be incorporated into FireCloud. Further cost and latency reductions may be possible through the adoption of these new technologies.

## Acknowledgments

This work was conducted and funded as part of the National Cancer Institute’s Cancer Genomics Cloud Pilots (HHSN261201400006C). We are grateful to Broad Institute’s Data Sciences Platform for collaborating to build and maintain FireCloud, and to our University of California collaborators, David Haussler and the Computational Genomics Laboratory at UC Santa Cruz and David Patterson and the AMPLab at UC Berkeley. Many thanks to Juli Klemm, Head of the Cancer Biology and Genomics Section, National Cancer Institute Center for Biomedical Informatics and Information Technology, for her support and guidance during FireCloud’s development.

## Notes

Conflicts of Interest: Dr. Philippakis is a Venture Partner at GV, a subsidiary of Alphabet Corporation

